# DNA-dependent RNA cleavage by the Natronobacterium gregoryi Argonaute

**DOI:** 10.1101/101923

**Authors:** Sunghyeok Ye, Taegeun Bae, Kyoungmi Kim, Omer Habib, Seung Hwan Lee, Yoon Young Kim, Kang-In Lee, Seokjoong Kim, Jin-Soo Kim

**Affiliations:** Center for Genome Engineering, Institute for Basic Science, Seoul 08826, South Korea; IBS School, University of Science and Technology, Daejeon 34113, South Korea; ToolGen, Inc., Seoul 08501, South Korea; Department of Chemistry, Seoul National University, Seoul 08826, South Korea

## Abstract

We show here that, unlike most other prokaryotic Argonaute (Ago) proteins, which are DNA-guided endonucleases, the *Natronobacterium gregoryi*-derived Ago (NgAgo) can function as a DNA-guided endoribonuclease, cleaving RNA, rather than DNA, in a targeted manner. The NgAgo protein, in complex with 5’-hydroxylated or 5’-phosphrylated oligodeoxyribonucleotides (ODNs) of variable lengths, split RNA targets into two or more fragments in vitro, suggesting its physiological role in bacteria and demonstrating a potential for degrading RNA molecules such as mRNA or lncRNA in eukaryotic cells in a targeted manner.

---

Argonaute (Ago) proteins are complexed with small DNA or RNA guides and cleave nucleic acid targets whose sequences are complementary with the guides^1^. Agos in eukaryotes bind to small interfering RNAs (siRNAs) or microRNAs (miRNAs) to suppress gene expression by degrading mRNAs in a targeted manner, a regulatory process known as RNA interference (RNAi). Agos in eubacteria and archaea are complexed with DNA or RNA guides and function as defense systems against foreign mobile genetic elements such as phages or plasmids^2, 3^.

Recently, Gao et al. reported that the Ago homolog found in *Natronobacterium gregoryi* (NgAgo) utilized 5’ phosphorylated small guide DNA molecules to recognize and cleave double-stranded DNA in vitro and in human cell lines, enabling DNA-guided genome editing^4^. Unlike CRISPR-Cas9 (clustered regularly interspaced short palindromic repeats (CRISPR) – CRISPR-associated protein 9)^5–8^ or Cpf1 (CRISPR from *Prevotella* and *Francisella* 1) systems ^9–12^, which are limited by the requirement of proto-spacer adjacent motif (PAM) sequences recognized by Cas9 or Cpf1, prokaryotic Agos in general and NgAgo in particular do not require a PAM, an important advantage of DNA-guided genome editing over RNA-guided editing with CRISPR systems.

Unfortunately, however, we and others failed to detect small insertions or deletions (indels) at NgAgo target sites in human cells^13, 14^, mice^13^, or zebrafish^15^, bona fide evidence of genome editing via non-homologous end-joining (NHEJ) repair of DNA double-strand breaks (DSBs) in eukaryotic cells. Nevertheless, several groups reported that NgAgo achieved DNA-guided gene knockdown in human cells^4, 13^ and zebrafish^15^, leading to confusion in the scientific community^16^. Here, we show that NgAgo utilizes both 5’-hydroxylated and 5’-phosphorylated guide oligodeoxyribonucleotides (gODNs) to cleave RNA, rather than DNA, in a sequence-dependent manner, suggesting its mechanism of action and potential applications in biomedical research.

First, we cloned the gene encoding NgAgo in an *E. coli* expression vector and purified the recombinant NgAgo protein fused to a poly-histidine tag by metal affinity chromatography (**Supplementary Methods and Supplementary Fig. 1**). The NgAgo protein was incubated with a 5’-hydroxylated or 5’-phosphorylated gODN at 37 ℃ for 30 minutes to form Ago deoxyribonucleoprotein (DNP) complexes. An RNA target (**Supplementary Table. 1**), prepared by in vitro transcription with T7 RNA polymerase, was treated with the NgAgo DNP at 37 ℃ for 20-60 minutes. NgAgo in complex with the antisense gODN whose sequence was complementary with the RNA target encoding exon 11 of the human DYRK1A gene cleaved the RNA transcript site-specifically (Fig. 1a). In contrast, NgAgo pre-incubated with a sense gODN did not cleave the target (Fig. 1b), suggesting that base pairing between RNA and gODN is essential for targeted RNA cleavage. Both 5’-hydroxylated and 5’-phosphrylated gODNs were able to direct NgAgo to cleave the RNA transcript. To the best of our knowledge, no other eukaryotic or prokaryotic Ago proteins are guided by 5’-hydroxylated gODNs. A small antisense RNA, the size and sequence of which was identical with those of the gODN, failed to guide NgAgo to cleave the RNA target in vitro (Fig. 1a). Single-stranded and double-stranded DNA molecules were not cleaved by the NgAgo DNP under our experimental conditions. Taken together, these results show that NgAgo is a DNA-guided ribonuclease (RNase) rather than a DNA-guided deoxyribonuclease (DNase).

**Figure 1.**
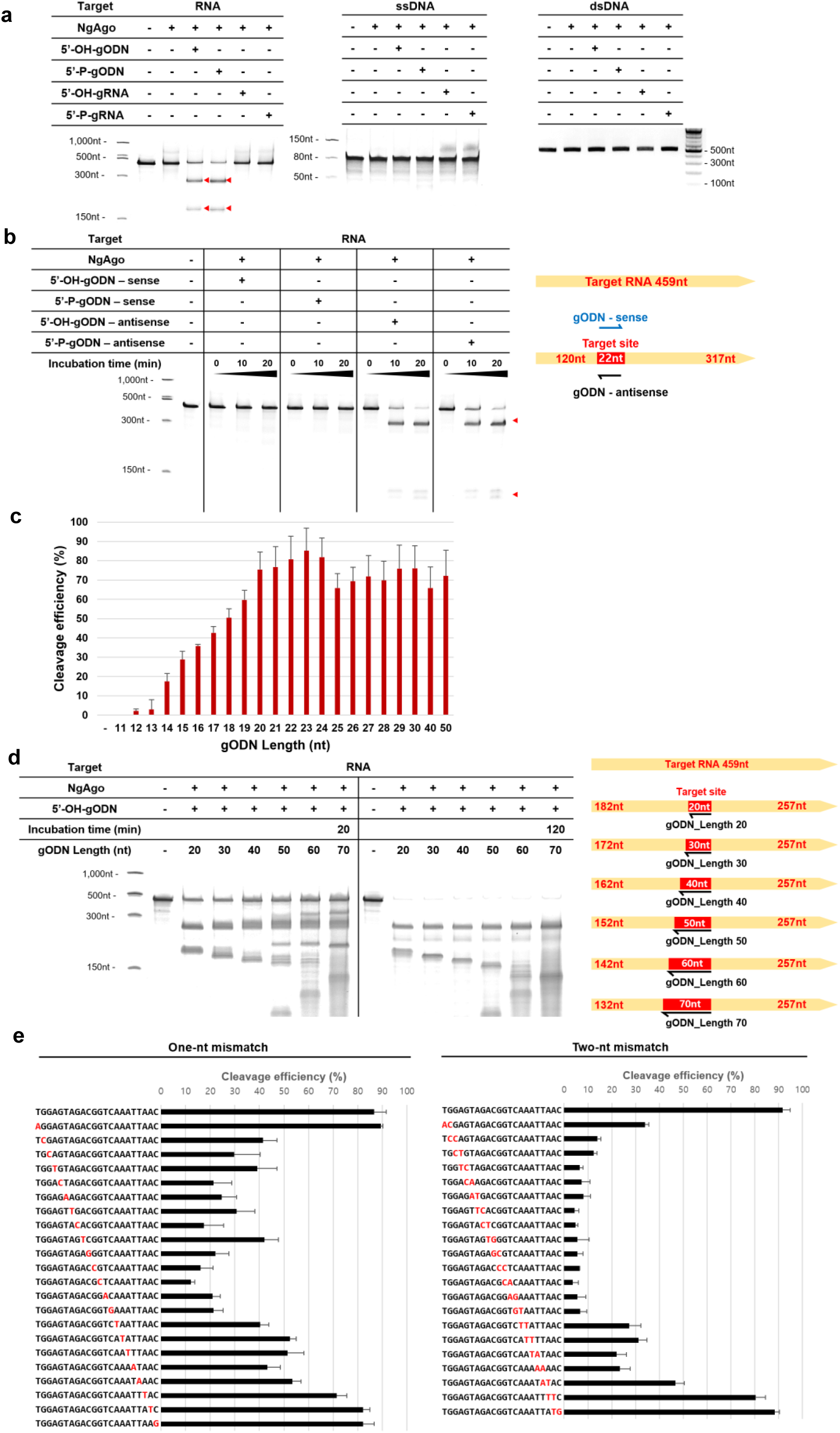
NgAgo-mediated RNA cleavage *in vitro.* (a) RNA, single-stranded DNA (ssDNA),and double-stranded DNA (dsDNA) targets were treated with the purified NgAgo protein in the presence of 5’-hydroxylated (5’-OH) or 5’-phosphorylated (5’-P), 22-nt guide ODNs (gODNs) or RNAs (gRNAs). (b) The RNA substrate was treated with NgAgo in the presence of gODNs in sense or antisense directions relative to the RNA target. Red triangles indicate positions of RNA fragments produced by NgAgo-mediated RNA cleavage in a denaturing urea-polyacrylamide gel. (c) RNA cleavage efficiencies of NgAgo with gODNs of variable lengths were measured in a urea-polyacrylamide gel. Error bars indicate s.e.m. (n = 3). (d) Urea-polyacrylamide gel showing intermediate RNA cleavage products. (e) Specificity of DNA-guided RNA cleavage by NgAgo. An RNA substrate was digested with NgAgo in the presence of a fully-matched or mismatched gODNs. Mismatched nucleotide in gODNs were shown in red. Error bars represent s.e.m. (n = 3).

We next investigated whether metal ions were required for RNA cleavage by NgAgo. In the presence of EDTA, a metal ion chelator, NgAgo failed to cleave the RNA target (**Supplementary Fig. 2a**). To identify optimal cations, we incubated the NgAgo DNP with a series of metal ions. Among 9 divalent or trivalent metal ions, Mn^++^ was most efficient at a concentration of 10 μM. The NgAgo DNP was also able to cleave RNA in the presence of Mg^++^ ions, albeit at a much higher concentration (≥ 100 μM). NgAgo failed to show > 10% RNA cleavage in the presence of all of the other cations we tested (**Supplementary Fig. 2b**). Accordingly, we performed NgAgo reactions in the presence of 10 μM Mn^++^ throughout this study unless indicated otherwise.

We also tested gODNs of variable lengths to optimize DNA-guided RNA cleavage by NgAgo (Fig. 1c). ODNs of ≤ 13 nucleotides (nt) in length failed to guide NgAgo to cleave RNA efficiently. The efficiency of RNA cleavage was proportional to the length of gODNs up to a point. The 22-nt gODN was as efficient as ≥ 23-nt gODNs. Therefore, we used 22-nt gODNs in this study unless indicated otherwise. To determine exact positions of RNA cleavage by NgAgo, we cloned and sequenced RNA products after poly(A) tailing (**Supplementary Fig. 3**). The RNA substrate was cut at several positions within a 22-nt gODN-complementary target region. We also noted that NgAgo in complex with ≥ 30-nt gODNs initially yielded RNA fragments distinct from those produced by NgAgo in complex with 20-nt gODNs (Fig. 1d). Upon complete digestion, most of these additional RNA products disappeared, suggesting that they were intermediates. These results show that an RNA substrate can be cut at multiple sites by NgAgo and that NgAgo can cleave the RNA strand in an extended DNA-RNA duplex, reminiscent of RNase H-like activity^17^. Note, however, that we purified the recombinant NgAgo protein to near homogeneity by two different methods (**Supplementary Methods**) to rule out the possibility that the DNA-dependent RNase activity we observed with NgAgo was caused by contaminating *E. coli* RNase H.

We also performed NgAgo reactions under multiple turnover conditions by incubating 0.03 μM NgAgo with 0.1 μM RNA substrate. NgAgo cleaved the RNA substrate completely in 16 hours (**Supplementary Fig. 4**). This result show that, unlike Cas9, a single-turnover enzyme^18^, NgAgo is a multiple-turnover enzyme, cleaving its substrate repeatedly.

To investigate whether NgAgo has off-target effects, we tested the protein premixed with a series of 22-nt gODNs with one– or two-nt mismatches (Fig. 1e). Most of these mismatched ODNs were associated with reduced RNA cleavage activities, compared to a fully-matched gODN. In particular, NgAgo complexed with ODNs harboring mismatches in nt positions 5-14 (numbered from 5’ to 3’) were poorly active, suggesting that these positions constitute a “seed region”. NgAgo tolerated mismatches at the 5’ and 3’ ends.

We next tested several antisense gODNs whose sequences were complementary with the DYRK1A exon 11 RNA transcript. All 7 NgAgo DNPs cleaved the DYRK1A transcript, producing RNA bands at expected positions in a denaturing urea-polyacrylamide gel (Fig. 2). Efficiencies of RNA cleavage were not very dependent on gODNs, suggesting that NgAgo does not discriminate target RNA sequences.

**Figure 2.**
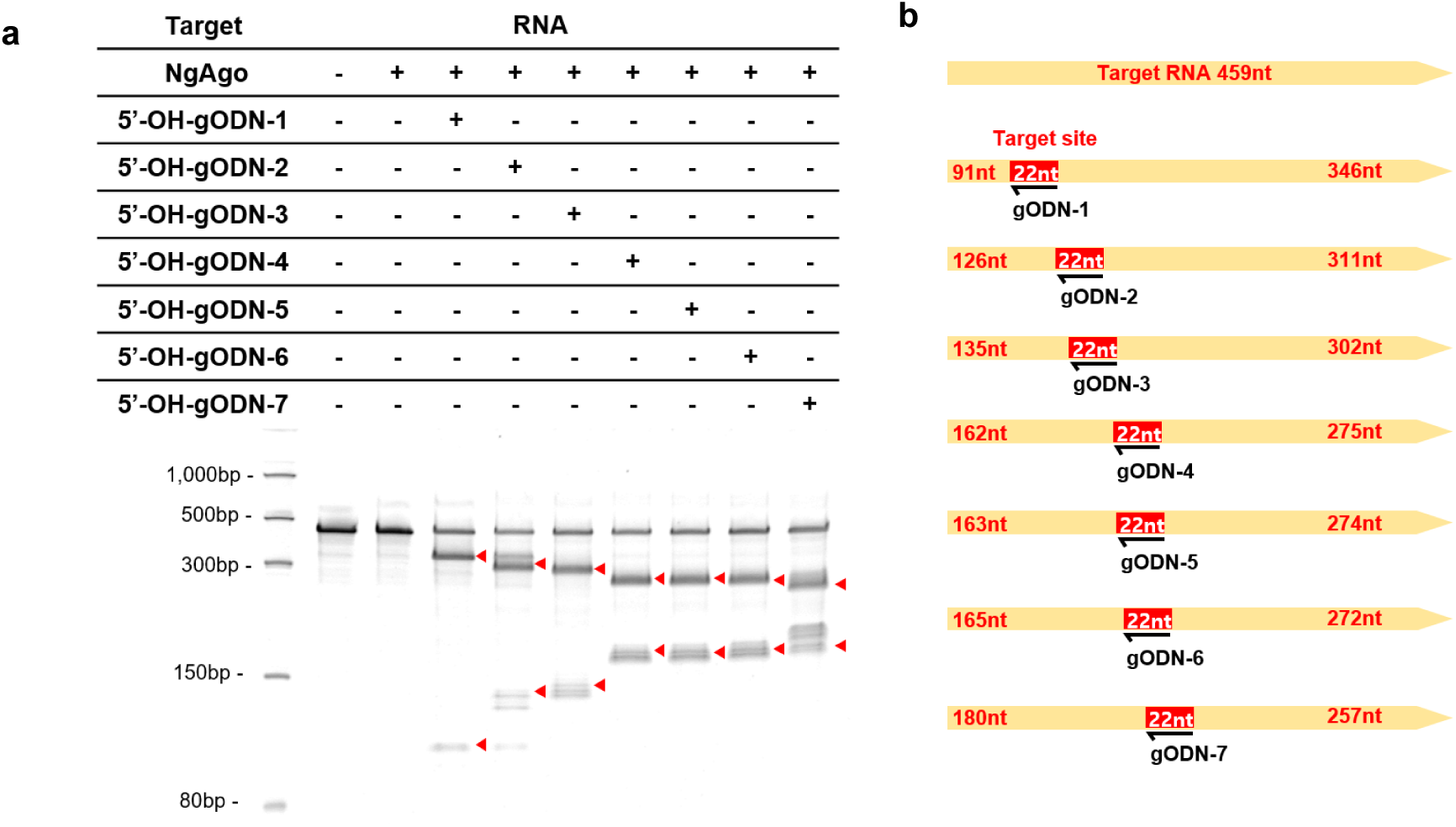
gODN sequence-dependent RNA cleavage by NgAgo. (a) An urea-polyacrylamide gel showing RNA cleavage by NgAgo with a series of antisense gODNs. Red arrow indicates cleavage products. (b) Schematic representation of the RNA substrate and gODNs targeted to different sites.

Several recent reports claimed that NgAgo-DNA complexes achieved knockdown of the green fluorescent protein (GFP) gene^4, 13, 16^ and endogenous genes^15^ in human cells and zebrafish, respectively. Our results suggest that NgAgo-mediated gene knockdown in cells can be at least partially attributed to the DNA-dependent RNase activity of NgAgo. Qi et al. hypothesized that NgAgo may bind to a target gene to block its transcription, because three NgAgo mutants (D663A, D738A, and D663A/D738A), presumably lacking catalytic activity, were still able to cause the same phenotype as wild-type NgAgo in zebrafish^15^. We found, however, that these and other mutants with an aspartate-to-alanine substitution cleaved an RNA target as well as wild-type NgAgo in vitro (**Supplementary Fig. 5**), suggesting that these residues do not constitute an active site and that the RNase activity of the NgAgo mutants was still responsible for the phenotype.

Taken together, our data show that NgAgo is guided by ODNs of variable lengths to split RNA into two or more fragments. We hypothesize that NgAgo and its orthologues^19^ protect host cells from mobile genetic elements such as bacteriophages by degrading RNA strands in RNA-DNA duplexes. We also propose that the DNA sequence-specific RNase activity of NgAgo DNPs can be harnessed for biomedical research, mediating the degradation of mRNA or functional RNAs such as long non-coding RNAs (lncRNAs) and circular RNAs in a targeted manner. Unlike siRNAs, NgAgo-gODN DNPs are orthogonal in eukaryotic cells: gODNs do not compete with endogenous miRNAs to complex with host Ago proteins, a potential advantage over RNAi.

## ACKNOWLEDGMETNS

This work was supported by IBS-R021-D1 (to J.-S. K.).

## REFERENCES

1. Swarts D.C. et al. Nature 507, 258–261 (2014).

2. Kaya E. et al. Proc. Nat. Acad. Sci. USA 113, 4057–4062 (2016).

3. Swarts D.C. et al. Nat. struct. Mol. Biol. 21, 743–753 (2014).

4. Gao F., Shen X.Z., Jiang F., Wu, Y. & Han, C. Nat. biotechnol. 34, 768–773 (2016).

5. Cho S.W., Kim S., Kim J.M. & Kim J.S. Nature biotechnol. 31, 230–232 (2013).

6. Cong L. et al. Science 339, 819–823 (2013).

7. Jinek M. et al. Science 337, 816–821 (2012).

8. Mali P. et al. Science 339, 823–826 (2013).

9. Hur J.K. et al. Nat. biotechnol. 34, 807–808 (2016).

10. Kim D. et al. Nat. biotechnol. 34, 863–868 (2016).

11. Kim Y. et al. Nat. biotechnol. 34, 808–810 (2016).

12. Zetsche B. et al. Cell 163, 759–771 (2015).

13. Burgess S. et al. Protein & cell (2016).

14. Lee S.H. et al. Nat. biotechnol. (2016).

15. Qi J. et al. Cell Res. (2016).

16. Cyranoski D. Nature 536, 136–137 (2016).

17. Donis-Keller H. Nucleic acids research 7, 179–192 (1979).

18. Sternberg S.H., Redding S., Jinek M., Greene E.C. & Doudna J.A. Nature 507, 62–67 (2014).

19. Yuan Y.R. et al. Molecular cell 19, 405–419 (2005).

